# Brown and Lesser noddies as epidemiological reservoirs and sentinels of avian influenza virus in the South-western Indian Ocean

**DOI:** 10.64898/2026.03.31.715511

**Authors:** Camille Lebarbenchon, Céline Toty, Nina Voogt, Christine Larose, Audrey Jaeger, Cheryl Sanchez, Sophie Bureau, Liadrine Moukendza-Koundi, Muriel Dietrich, Nirmal Shah, Chris Feare, Byron Göpper, Matthieu Le Corre, Karen D. McCoy

## Abstract

Avian influenza virus (AIV) epidemiology is well-documented in temperate regions but remains poorly understood in isolated ecosystems like tropical oceanic islands. On these islands, seabirds nest in dense interspecific colonies where the role of different species as reservoirs and dispersers of AIV may vary greatly. Here, we examine the role of noddies (*Anous spp*.) as potential reservoirs for low pathogenic AIV and evaluate their potential as sentinel species for highly pathogenic AIV introduction on tropical oceanic islands. We analyzed blood samples from 11 seabird species across eight islands in the southwestern Indian Ocean (2015–2020). Noddies exhibited high, stable seroprevalence (30–45%), comparable to reservoir host species in temperate regions. The detection of two N7-positive noddies, sampled the same year on two distinct islands, provided direct molecular evidence that AIV actively circulates on these island colonies. While most other species showed low exposure, Bridled Terns (*Onychoprion anaethetus*) had exceptionally high seroprevalence (80%), though their reservoir status requires further investigation due to limited sampling. Given noddies’ consistent exposure and regional distribution, we recommend prioritizing islands with large noddy populations for AIV surveillance. Continued investigation of viral dynamics within and among islands is now called for to elucidate the ecological drivers of AIV maintenance and transmission.

---

The epidemiology of avian influenza viruses (AIV) has been extensively studied in temperate regions of the Northern Hemisphere, however, significant knowledge gaps persist in more isolated ecosystems, such as oceanic islands. These islands host avian communities dominated by seabirds and vagrant shorebirds, with a notable absence of native Anseriformes, the primary reservoir hosts of AIV in the temperate Northern Hemisphere regions (e.g. ducks and geese; [1–3]). The distinct community composition found on oceanic islands may constrain AIV transmission patterns and limit viral diversity to subtypes typically associated with marine avian hosts (e.g. H13–H16; [4–7]). Additionally, the ecology of seabirds, characterized by prolonged pelagic phases and seasonal breeding in dense colonies, may further influence the spatial and temporal dynamics of viral transmission. These unique conditions raise critical questions about how AIV circulates and persists in these ecologically distinct ecosystems.

The southwestern Indian Ocean hosts some of the world’s largest seabird breeding colonies, with an estimated breeding population of 19 million individuals [8]. These islands are critical breeding sites for several seabird taxa, including Charadriiformes (terns), Phaethontiformes (tropicbirds), Procellariiformes (petrels and shearwaters), and Suliformes (boobies and frigatebirds), which often aggregate in exceptionally dense colonies of hundreds of thousands to millions of birds. Most seabirds in this region are highly pelagic, without contact with continental landmasses. For instance, adult Sooty Terns (*Onychoprion fuscatus*) undertakes extensive migrations between breeding seasons (covering distances >50,000 km; [9]) as well as juveniles during the < 6 years they take to attain maturity. They, however, remain almost entirely at sea, potentially limiting opportunities for virus transmission. In contrast, Brown noddies (*Anous stolidus*) and Lesser noddies (*Anous tenuirostris*) exhibit movement patterns that could actively facilitate virus dispersal across the oceanic basin [10]. Tracking studies have demonstrated that these species frequently move between islands during the non-breeding season, roosting on multiple islands from the Tanzanian coasts to the Maldives archipelago, thereby creating high connectivity within the western Indian Ocean [10]. Furthermore, the detection of AIV on Réunion Island, phylogenetically related to viruses circulating in the Northern Hemisphere, demonstrates that inter-hemispheric viral exchange occurs, challenging the assumption that small tropical islands could be protected from AIV introductions [11]. Indeed, it was recently shown that AIV strains (H5N1 virus clade 2.3.4.4b) implicated in the outbreak on the sub-Antarctic archipelagos of Crozet and Kerguelen originated from the distant South Georgia islands in the Southern Atlantic, and that multiple introductions likely occurred [12]. Together, these observations underscore the need to investigate how AIV circulate within and between apparently isolated, yet ecologically interconnected, island systems.

Seabird community composition and abundance exhibit marked spatial variability across tropical islands, with each island hosting a distinct assemblage of breeding species. Within this heterogeneous system, Brown and Lesser noddies may play a central epidemiological role. Their colonial breeding behavior and flexible movement strategies, ranging from year-round residency to long-distance movements between islands, could facilitate both local persistence and regional dispersal of AIV [10]. This combination of connectivity and site fidelity highlights the potential role of noddies as potential keystone species in island AIV dynamics. In the context of the global spread of highly pathogenic (HP; e.g. [5–7,12,13]) H5 viruses, these traits could also make noddies valuable sentinel species for epidemiological surveillance in small island ecosystems. Identifying species occupying such a central role in these spatially structured avian communities is essential for the development of targeted, island-specific, surveillance programs.

In this study, we investigated the potential role of Brown and Lesser noddies as epidemiological reservoirs for AIV across their geographic range in the southwestern Indian Ocean. We hypothesized that, if these species act as reservoirs, their seroprevalence would remain relatively high and stable across islands and years, reflecting their extensive movement patterns and regional population connectivity. To provide a broader epidemiological context, we also assessed AIV exposure in three tern species, Sooty Terns, Bridled Terns (*Onychoprion anaethetus*), and White Terns (*Gygis alba*), that breed sympatrically on the same islands as noddies. Additionally, we included other seabird species to confirm their limited role in AIV epidemiology compared to noddies and terns (shearwaters, tropicbirds, boobies and frigatebirds). We discuss the dual role of noddies as both potential reservoirs and sentinel species for low pathogenic (LP) AIV surveillance, emphasizing their importance for early detection systems in the event of HP AIV epizootics on tropical oceanic islands.

## Material and methods

### Study sites

We focused on eight islands in the Western Indian Ocean: Bird (3°43’S, 55°12’E), Cousin (4°19’S, 55°39’E), Cousine (5°39’S, 55°39’E), Europa (22°21’S, 40°21’E), Juan de Nova (17°03’S, 42°45’E), Lys (Glorieuses archipelago, 12°29’S, 47°23’E), Réunion (21°22’S, 55°34’E), and Tromelin (15°53’S, 54°31’E). These islands host highly heterogeneous seabird communities in terms of species richness, population size, and density [14–16], and represent critical breeding grounds for tropical seabirds in the Indian Ocean [17]. They were selected to cover the broadest possible geographic range of Lesser and Brown noddies in the region (Figure 1). While Brown and Lesser noddies do not breed on Juan de Nova and Europa, we included these islands to assess AIV circulation in their absence. These islands host massive breeding colonies of Sooty Terns [8,14], as well as other abundant seabird species, particularly on Europa, including Red-footed Boobies (*Sula sula*), White-tailed Tropicbirds (*Phaethon lepturus*), and Great Frigatebirds (*Fregata minor*). Réunion Island hosts only a small breeding population of Brown noddies, but thousands of both Brown and Lesser noddies roost year-round, probably originating from other nearby colonies.

**Figure 1.**
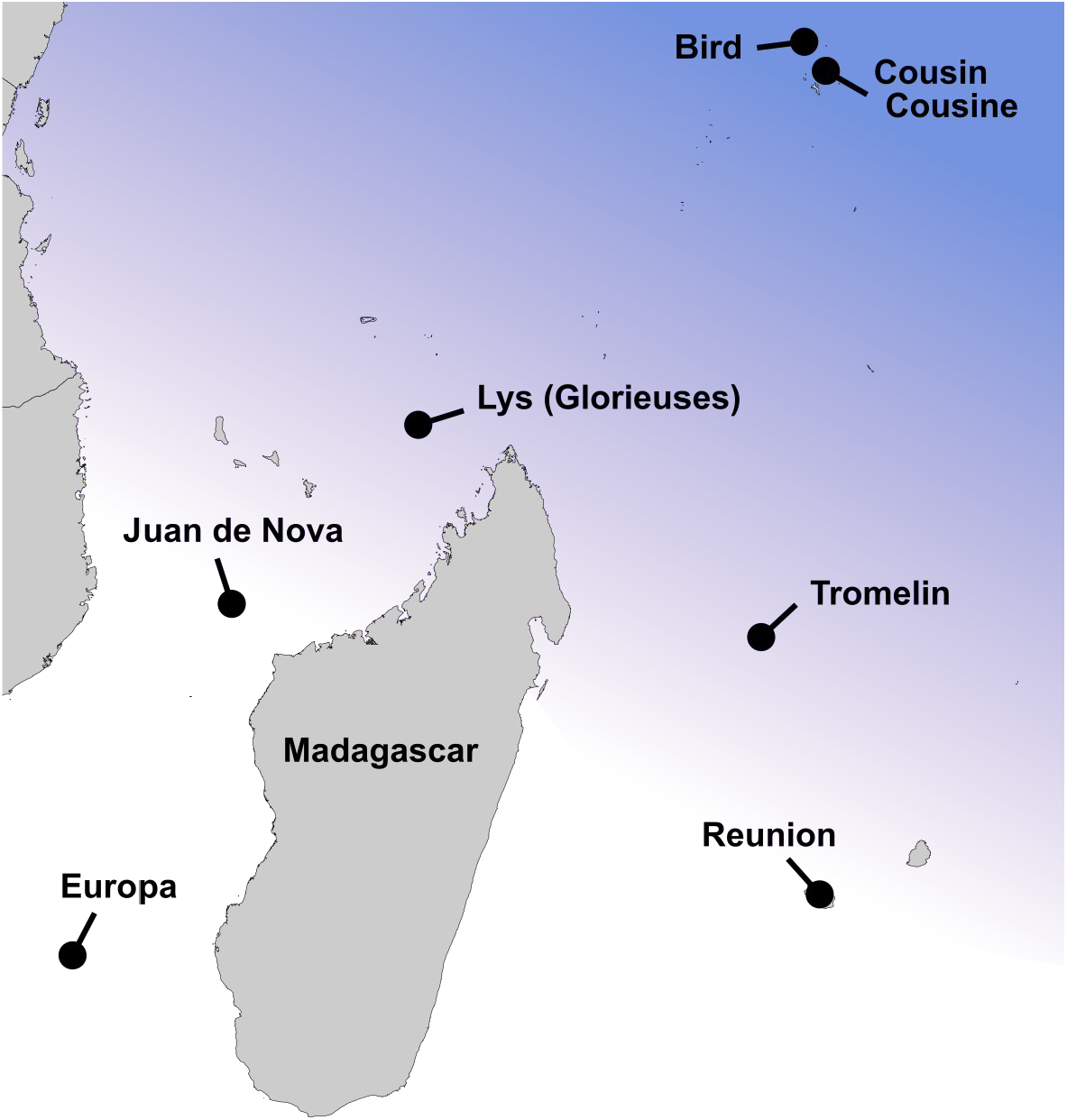
Study sites and distribution range of Brown noddies (*Anous stolidus*) and Lesser noddies (*Anous tenuirostris*) in the southwestern Indian Ocean. The shaded blue area represents the geographic distribution of both species (adapted from [18]). Islands included in this study are indicated by black dots (due to their geographic proximity Cousin and Cousine are combined in a single dot).

### Ethic statement and research permits

All procedures were evaluated and approved by the Ethics Committee of La Réunion (agreement number A974001) and authorized by the French Ministry of Education and Research (APAFIS#3719-2016012110233597v2). Bird capture, handling, and collection of biological material were approved by the Center for Research on Bird Population Biology (PP616; National Museum of Natural History, Paris, France) for Europa, Juan de Nova, Lys (Glorieuses), Réunion, and Tromelin Islands, and by the Seychelles Bureau of Standards and the Seychelles Ministry of Agriculture, Climate Change and Environment for Bird, Cousin, and Cousine Islands (Mahe, Seychelles).

### Bird sampling

Samples were collected between 2015 and 2020 to investigate species-related and temporal variation in AIV antibodies and viral shedding (Table 1). A small blood sample (up to 1.0% of body weight) was collected from the medial metatarsal vein and centrifuged within 3 hours of collection. Sera were stored at −20°C until testing. Faecal (cloacal swab) and saliva (oropharyngeal swab) were collected using sterile rayon-tipped applicators (Puritan, USA). Both swabs were placed in the same tube containing 1 ml of virus transport media, composed of brain heart infusion (Conda, Spain) supplemented with penicillin G (1000 units/ml), streptomycin (1 mg/ml), kanamycin (0.5 mg/ml), gentamicin (0.25 mg/ml), and amphotericin B (0.025 mg/ml) [19]. Due to the lack of cold storage facilities on Lys Island, swabs were placed in RNAprotect (QIAGEN, USA) instead of virus transport media, and were stored in a cooler box with ice packs for three days in the field. Plasma and swabs were stored at −20°C and −80°C, respectively, in the laboratory until testing.

**Table 1.**
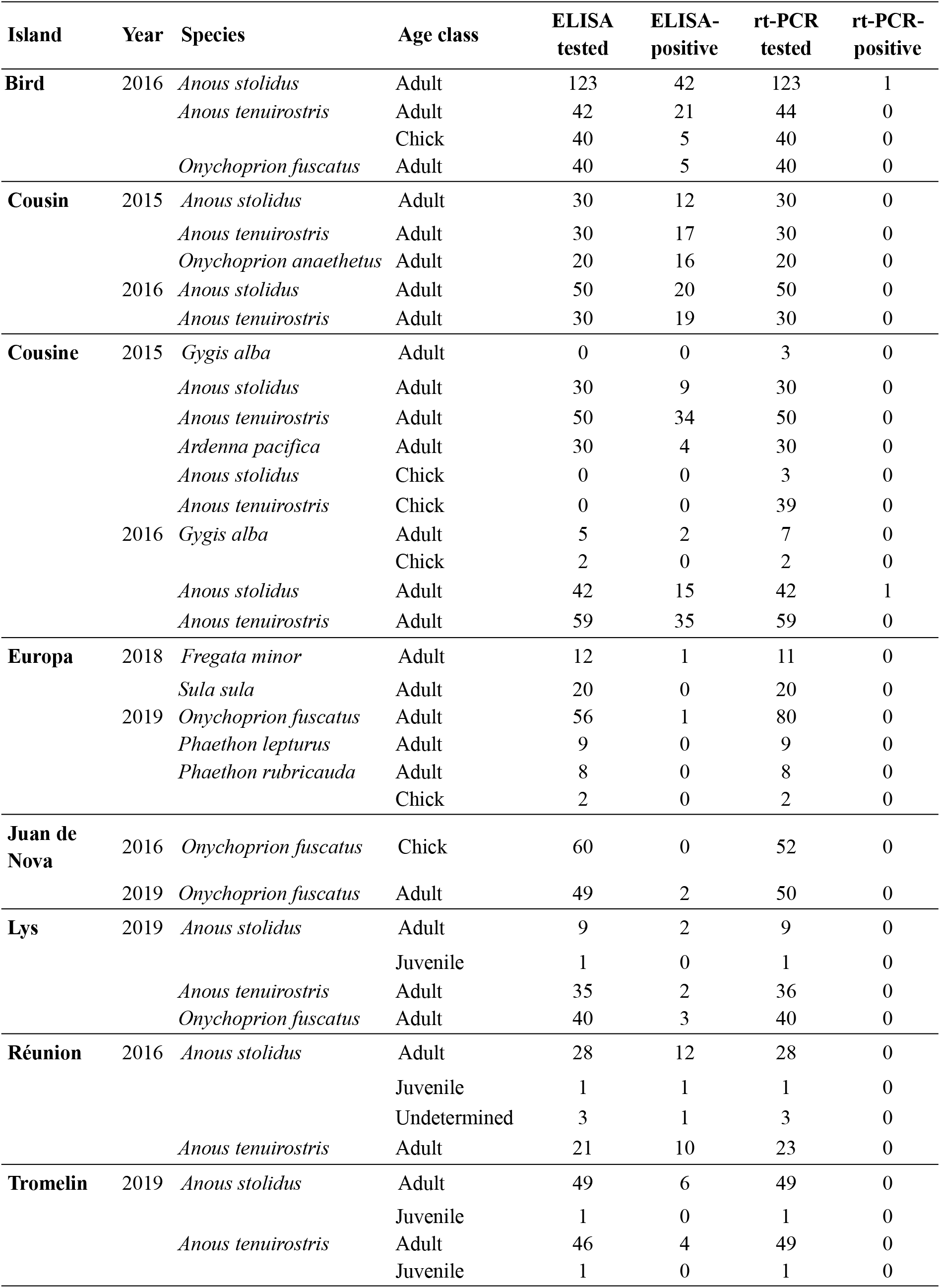
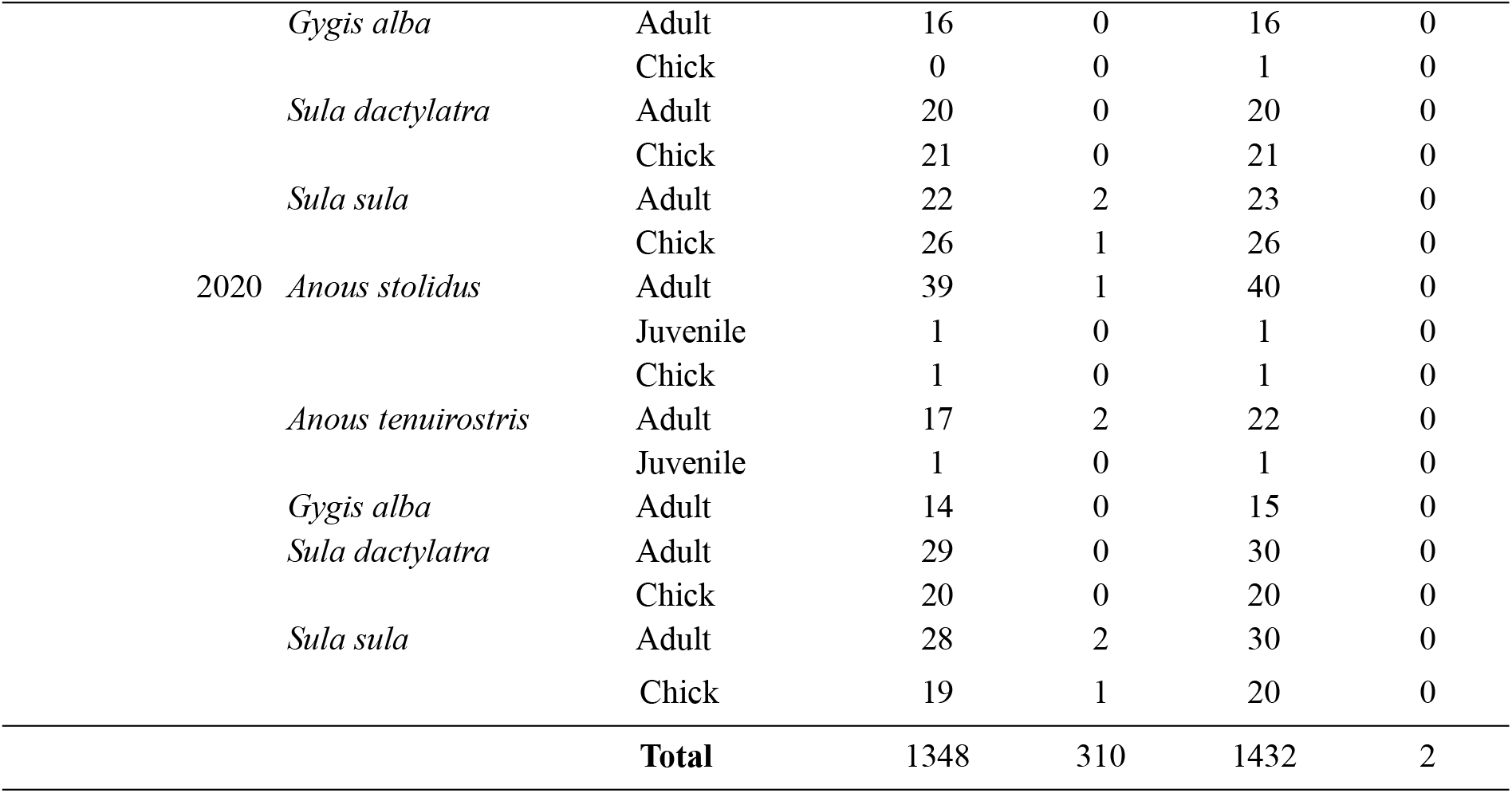
Origin and number of tested samples with the enzyme-linked immunosorbent assay (ELISA) and the real-time polymerase chain reaction (rt-PCR).

### Serology

Sera were tested using the IDvet ID Screen Influenza A Antibody Competition enzyme-linked immunosorbent assay (ELISA; IDvet, France), following an optimized protocol for the detection of AIV-specific nucleoprotein (NP) antibodies in wild birds [20]. This protocol was previously used to detect seropositive seabirds in the Western Indian Ocean [10,11]. Sample absorbance was measured at 450 nm using a Sunrise microplate reader (TECAN, Austria). Samples with a sample-to-negative control ratio (S/N) below 0.4 were considered positive for AIV NP antibodies, while samples with S/N ≥ 0.55 were considered negative. Samples yielding S/N values between 0.4 and 0.55 were retested; based on the second S/N result, they were classified as negative (S/N > 0.4) or positive (S/N ≤ 0.4).

We analyzed the data using Generalized Linear Models (GLMs) with a binomial error distribution and logit link function, to assess the effects of bird species, island, age class (chick, juvenile, adult) and their interactions, on the probability of detecting AIV NP antibodies in plasma samples. Due to the significantly higher seropositivity in adult birds, we also analyzed the effects of species and island on the probability of detecting AIV NP antibodies specifically in this age class. For Bird Island, previously published data were included to investigate inter-annual variation in AIV NP antibodies in Lesser noddies, Brown noddies, and Sooty terns (additional data: N = 1,244 adult birds from [10,11]). Analyses were conducted in R 4.5.1 [21].

### Molecular detection

Swab were thawed overnight at 4°C, briefly vortexed, and centrifuged at 1,500g for 15 minutes. Nucleic acids were extracted using the IndiSpin QIAcube HT Pathogen Kit (QIAGEN, USA). Reverse transcription was performed on 10μl of RNA using ProtoScript II Reverse Transcriptase and Random Primer 6 (New England BioLabs, USA), as previously described [10]. Complementary DNA (cDNA) was then tested for the presence of the AIV Matrix (M) gene by real-time polymerase chain reaction (rt-PCR) [22], using a CFX96 Touch Real-Time PCR Detection System (Bio-Rad, USA). Subtyping was attempted using primer sets designed to amplify avian HA (H1–H16) and NA (N1–N9) subtypes of AIV [22–26]. Amplification products were analyzed by electrophoresis on a 2% agarose gel stained with GelRed (Biotium, USA). Amplicons were sequenced by Genoscreen (Lille, France).

## Results

### Seroprevalence

A total of 1,348 plasma samples were collected from 11 seabird species across eight islands in the southwestern Indian Ocean. Overall, 310 samples (23.0%) tested positive for AIV NP antibodies (Table 1). Significant differences in seropositivity were detected among species (χ^2^ = 57.5; df = 10; p < 0.001), among islands (χ^2^ = 315.8; df = 7; p < 0.001), and among age classes (χ^2^ = 15.8; df = 2; p < 0.001). Due to unbalanced sampling between age classes (N = 1,148 adults *vs* N = 200 for chicks and juveniles combined) and the limited number of seropositive chicks and juveniles (N = 8) compared to adults (N = 301), only adult birds were included in subsequent analyses.

Significant differences in seroprevalence were detected among adult seabird species (χ^2^ = 61.6; df = 10; p < 0.001; Figure 2), with the highest prevalences observed in Bridled terns (80.0 ± 17.5%), followed by Lesser noddies (44.6 ± 5.3%), Brown noddies (29.8 ± 4.5%), Wedge-tailed shearwaters (13.3 ± 12.1%), Great frigatebirds (*Fregata minor*; 8.3 ± 15.6%), Sooty terns (6.0 ± 3.4%), White terns (*Gygis alba*; 5.7 ± 7.7%), and Red-footed boobies (*Sula sula*; 5.7 ± 5.4%). No AIV NP antibodies were detected in White-tailed tropicbirds (*Phaethon lepturus*), Red-tailed tropicbirds (*Phaethon rubricauda*), or Masked boobies (*Sula dactylatra*).

**Figure 2.**
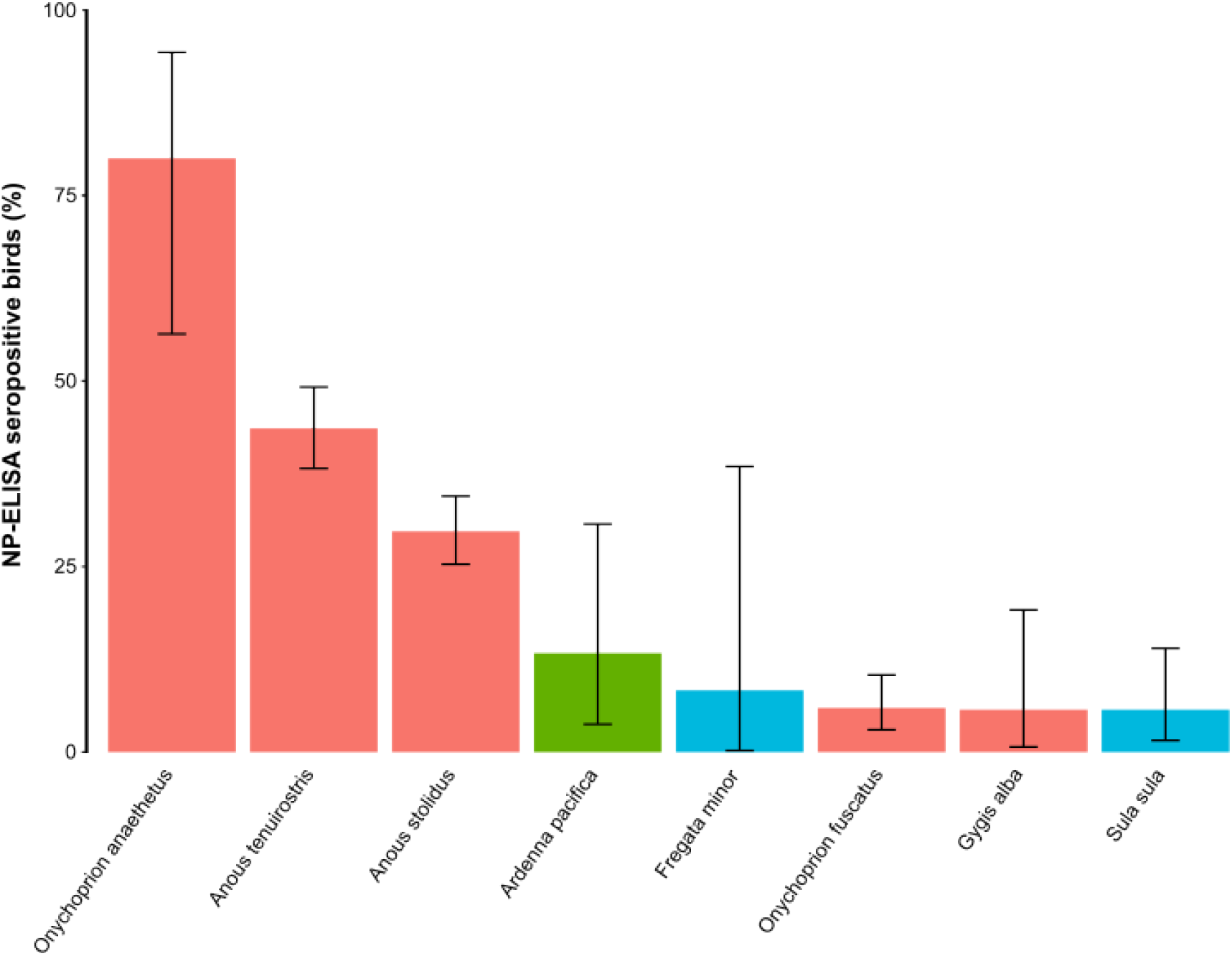
Seroprevalence of avian influenza virus nucleoprotein (NP) antibodies in seabirds from the Western Indian Ocean. Colors indicate bird orders: red, Charadriiformes; green, Procellariiformes; blue, Suliformes. Error bars represent 95% confidence intervals. As all tested individuals of White-tailed and Red-tailed tropicbirds (Phaethontiformes), and Masked boobies (Suliformes) were negative, these species were excluded from the figure.

Seroprevalence also varied significantly among islands (χ^2^ = 260.8; df = 7; p < 0.001), likely reflecting the uneven distribution of species. Islands hosting Bridled Terns, Lesser Noddies, or Brown Noddies, exhibited markedly higher seroprevalence (*e*.*g*. Cousin: 52.5 ± 7.7%; Cousine: 45.8 ± 6.6%; Bird: 33.2 ± 6.5%) than islands without these species. For example, Europa, where breeding species include Sooty Terns, White-tailed tropicbirds, Red-tailed tropicbirds, Red-footed boobies, and Great frigatebirds, had a very low overall seroprevalence (1.9 ± 2.6%; Table 1).

We further examined inter-island variation in seroprevalence for species sampled on at least three islands: Lesser noddies, Brown noddies, and Sooty terns (Figure 3). Significant differences were detected among islands for all three species: Lesser Noddy (χ^2^ = 89.9; df = 5; p < 0.001), Brown Noddy (χ^2^ = 35.6; df = 5; p < 0.001), and Sooty Tern (χ^2^ = 8.3; df = 3; p < 0.05). Finally, we investigated inter-annual variation in seroprevalence on Bird Island for Lesser noddies, Brown noddies, and Sooty terns. Overall, significant differences were detected among species (χ^2^ = 252; df = 2; p < 0.001) and across years (χ^2^ = 33; df = 4; p < 0.001), but when each species was considered individually (Figure 4), inter-annual variation was not statistically significant: Lesser Noddy (χ^2^ = 5.9; df = 5; p = 0.20), Brown Noddy (χ^2^ = 7.6; df = 4; p = 0.11), and Sooty Tern (χ^2^ = 2.0; df = 4; p = 0.73).

**Figure 3.**
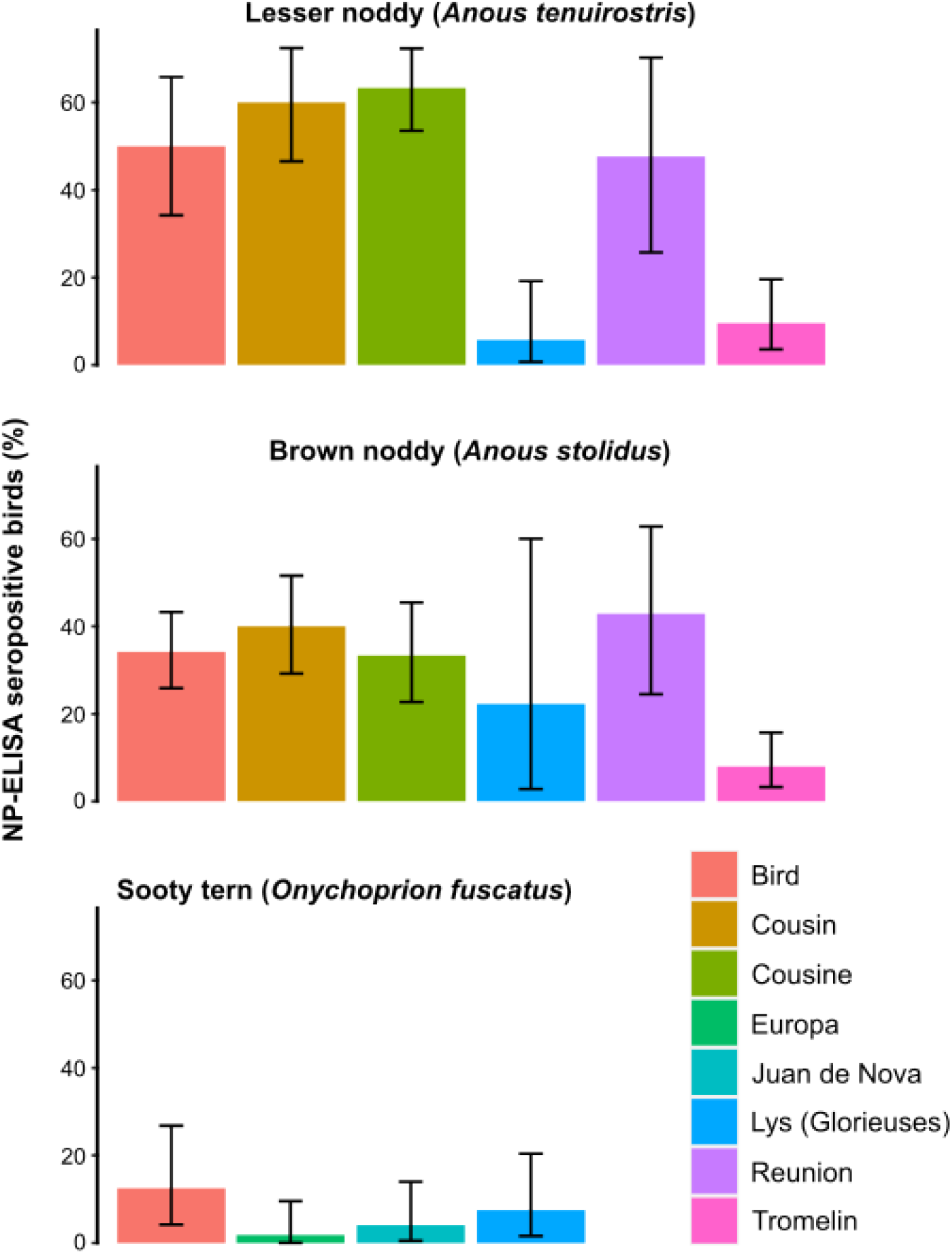
Inter-island variation of avian influenza virus nucleoprotein (NP) antibodies in Lesser noddies (*Anous tenuirostris*), Brown noddies (*Anous stolidus*), and Sooty Terns (*Onychoprion fuscatus*). Seroprevalence (with 95% confidence intervals) are presented for each species across multiple islands.

**Figure 4.**
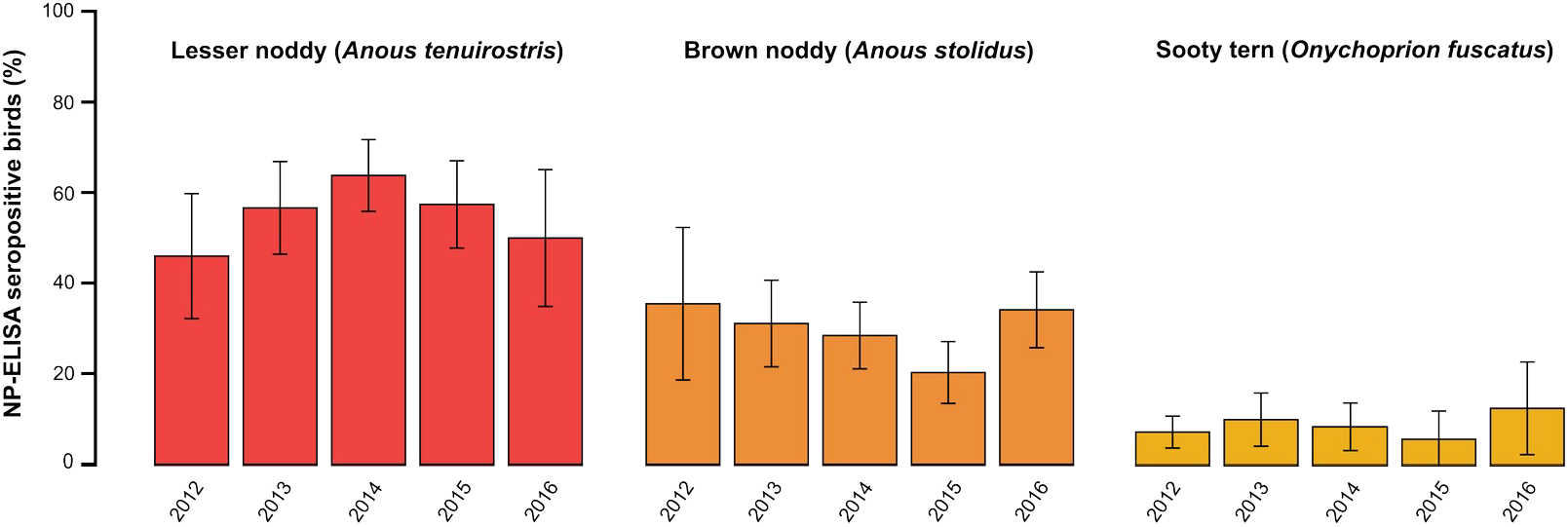
Inter-annual variation of avian influenza virus nucleoprotein (NP) antibodies in Lesser noddies (*Anous tenuirostris*), Brown noddies (*Anous stolidus*), and Sooty Terns (*Onychoprion fuscatus*), on Bird Island (2012-2016). Seroprevalence percentages (with 95% confidence intervals) are shown for each species over five consecutive years. Colors represent different species: Lesser noddies (red), Brown noddies (orange), and Sooty terns (yellow). 2012-2015 prevalence data were previously reported [10,11].

### Molecular detection

A total of 1,432 cloacal and oropharyngeal swabs were collected from seabirds (Table 1). Only two samples tested positive for the AIV M gene by rt-PCR. These positive samples were obtained from Brown noddies in 2016, on Bird and Cousine Islands. PCR assays targeting all 16 avian HA subtypes failed to identify the HA subtype in these two samples (i.e. all HA subtypes were tested by PCR, including H5 and H7, but none yielded positive results). However, both samples tested positive for the N7 NA subtype. Sequence analysis of the PCR amplicons revealed 99% nucleotide sequence identity (based on a 218 bp alignment). BLASTN 2.17.0+ analysis [27] revealed that the two N7 NA sequences shared >98% nucleotide identity with H10N7 viruses isolated from Northern Pintail (*Anas acuta*) and Northern Shoveler (*Spatula clypeata*) in Egypt, in 2012 (GenBank accession numbers KR862402 to KR862405). However, due to the limited length of the sequenced amplicons, interpretations regarding the host and geographic origin of these viruses should be made with caution.

## Discussion

This study provides compelling evidence that Brown and Lesser Noddies act as epidemiological reservoirs for AIV in the southwestern Indian Ocean. The high seroprevalence observed in these species (30-45%) are comparable to those reported in well-established reservoir hosts, such as ducks and gulls in temperate regions (e.g. [28]). The relative consistency in seroprevalence across years and among islands suggests that AIV exposure in noddies does not stem from sporadic epidemics which leave transient serological signatures. Instead, it likely reflects persistent viral circulation within these populations. Despite the limited genetic data available, the detection of two N7-positive Brown Noddies, sampled the same year on two distinct islands, provides direct molecular evidence that AIV actively circulates in seabird breeding colonies. These results build upon our earlier work during the non-breeding season where similarly high AIV exposure was documented and LP AIV were detected in noddies [10,11]. Collectively this work underscores the pivotal role of these species in the maintenance and regional dissemination of low-pathogenic AIV in tropical oceanic islands.

Beyond the role of Brown and Lesser noddies as reservoirs, this study highlights three key findings regarding interspecific variation in AIV exposure. First, we report high seroprevalence in Bridled Terns, with 80% of sampled individuals positive for AIV antibodies on Cousin Island. To our knowledge, very few studies have investigated AIV infection in this species, making this finding particularly notable [29,30]. While these data suggest substantial viral circulation, it remains unclear whether they reflect a single epidemic event within the studied population or rather that Bridled Terns like noddies act as epidemiological reservoirs for LP AIV. Further longitudinal sampling is now required to distinguish between these alternative hypotheses. Second, our results demonstrate that Wedge-tailed shearwaters experience regular AIV exposure, albeit with lower seroprevalence (13%). The consistent detection of seropositive individuals raises the hypothesis that shearwaters may acquire infections on islands where they co-occur with noddies, possibly through shared breeding habitats or foraging areas. Finally, we provide novel serological data for white terns, a species for which AIV exposure data were previously scarce [29]. The observed seroprevalence is comparable to that of Sooty terns (6%), though interpretations are limited by the relatively small sample size (N=37). Regarding other seabird groups, our findings confirm sporadic AIV exposure in boobies and frigatebirds, while indicating an absence of detectable viral circulation in tropicbirds [11].

To better elucidate the epidemiology of AIV in tropical island systems, future research should prioritize understudied species and taxonomic groups that may contribute to viral dispersal between islands or facilitate local maintenance of virus transmission. Among Charadriiformes, Caspian Terns (*Hydroprogne caspia*), Crested Terns (*Thalasseus bergii*), and Lesser-crested Terns (*Thalasseus bengalensis*) warrant particular attention. Their predominantly coastal distribution, in contrast to more pelagic species such as Sooty Terns and noddies, means that they could facilitate a transmission pathway, connecting the African continent, Madagascar, and islands of the southwestern Indian Ocean. Similarly, shorebirds, despite their limited population sizes [31,32], may contribute to long-distance viral dispersal owing to their migratory connectivity with Asia. Although their role in local AIV maintenance is likely limited, these species could nonetheless act as bridges for viral introduction between distant regions.

The potential role of landbird species, indigenous to tropical islands, remains largely unexplored. While sustained AIV transmission among terrestrial birds is unlikely, given their ecological separation from seabirds and their limited reservoir competence, spillover from domestic poultry represents a plausible source of exposure. Our observations of very low AIV exposure in both native and exotic landbird species on Réunion Island support this interpretation, suggesting that viral circulation in these species is more likely associated with anthropogenic interfaces (e.g., poultry farming) than with natural transmission cycles [33]. Further investigation of these dynamics would help elucidate island-specific ecological drivers of AIV maintenance and transmission.

Our findings provide a robust framework for designing island-based surveillance strategies aimed at monitoring both LP AIV circulation and the risk of H5N1 HP AIV introductions. Islands hosting breeding or roosting populations of Brown or Lesser Noddies should be prioritized, given their consistent role in viral maintenance. While seroprevalence could vary among islands, this variation might correlate with local noddy population sizes (e.g., lower on Tromelin and Lys, higher on Cousin and Cousine), with no evidence of entirely seronegative noddy populations. Additionally, islands with large Sooty Tern colonies, though less exposed to LP AIV, should also be targeted due to the potential for high-density transmission in the event of an HP AIV incursion, which could have devastating conservation implications.

Noddies fulfill nearly all criteria for effective sentinel species in AIV surveillance: high abundance, dense and accessible breeding colonies, consistent seroprevalence across seasons, and regional-scale movements that may facilitate early detection of viral introductions. One outstanding question is their susceptibility to HP AIV, for which no experimental or field data currently exist. In other tern and in gull species, HP H5N1 infections have caused severe clinical signs and mortality, demonstrating that some seabirds can serve as effective early-warning indicators (e.g. [34–36]). Determining whether noddies share this susceptibility is critical in order to complete the evidence base needed to formally recognize their sentinel role and strengthen early-warning surveillance capacity in small island states and territories.

## Acknowledgments

We thank Mickaël Baumann, Solenn Boucher, Mark Brown, Martin Cagnato, Emily Cavill, Quentin d’Orchymont, Alexis Cuvillier, Jérome Fort, Lucie Gauchet, Julien Gazal, David Gremillet, Yusuke Goto, Nicolas Guillerault, Jean Hivert, Marc Lemenager, Jade Lopez, Cédric Marteau, Erwann Moreau, Aine Nicholson, Aurélien Prudor, Merlène Saunier, for their help in bird capture and sampling. We are very grateful to the Savy family for their support in the fieldwork and sample collection on Bird Island, and to Mr Keeley the opportunity to conduct research on Cousine Island. We thank Eric Blais, Tom Hiney and Sam Hope, for organizing transportation and providing accommodation on Cousin Island, as well as to the volunteers for their warm welcome. Léon Biscornet, Graham Govinden and Brigitte Pool are also thanked for providing support for sample conservation at the Victoria Hospital (Mahe), and Helena Haber for arranging shipment of biological material to Réunion Island. Finally, we thank Enzo Moraes, Marie-Alice Simbi and Magali Turpin for providing technical support in the laboratory.

## Ethical standards

All procedures were evaluated and approved by the Ethics Committee of La Réunion (agreement number A974001) and authorized by the French Ministry of Education and Research (APAFIS#3719-2016012110233597v2).

## Data availability

Nucleotide sequences are available in Genbank under accession numbers PZ225334 and PZ225335.

## Declaration of AI use

Linguistic revision and editing of this manuscript were assisted by Le Chat (Mistral AI, Pro version, December 2025).

## Authors’ contributions

Conceptualisation : Camille Lebarbenchon.

Data Curation : Camille Lebarbenchon.

Formal Analysis : Camille Lebarbenchon, Muriel Dietrich.

Funding acquisition : Camille Lebarbenchon, Karen McCoy, Matthieu Le Corre.

Investigation : Camille Lebarbenchon, Nina Voogt, Byron Göpper, Sophie Bureau, Christine

Larose, Chris Feare, Audrey Jaeger, Matthieu Le Corre, Céline Toty, Karen McCoy, Muriel

Dietrich, Liadrine Moukendza-Koundi, Cheryl Sanchez

Project administration : Camille Lebarbenchon, Matthieu Le Corre, Nina Voogt, Byron

Göpper, Chris Feare, Nirmal Shah, Cheryl Sanchez.

Visualization : Camille Lebarbenchon, Muriel Dietrich.

Writing – original draft : Camille Lebarbenchon, Karen D. McCoy.

Writing – review & editing : All authors.

## References

1. Webster RG, et al. Evolution and ecology of influenza A viruses. Microbiological Reviews 1992; 56: 152–179.

2. Stallknecht DE, Shane SM. Host range of avian influenza virus in free-living birds. Veterinary Research Communications 1988; 12: 125–141.

3. Olsen B, et al. Global patterns of influenza A virus in wild birds. Science 2006; 312: 384– 388.

4. Verhagen JH, et al. Phylogeography and antigenic diversity of low-pathogenic avian influenza H13 and H16 viruses. Parrish CR, ed. Journal of Virology 2020; 94: e00537–20.

5. Wille M, et al. Extensive geographic mosaicism in avian influenza viruses from gulls in the Northern hemisphere. Davis T, ed. PLoS ONE 2011; 6: e20664.

6. Fouchier RAM, et al. Characterization of a novel influenza A virus hemagglutinin subtype (H16) obtained from Black-Headed Gulls. Journal of Virology 2005; 79: 2814–2822.

7. Kawaoka Y, et al. Is the gene pool of influenza viruses in shorebirds and gulls different from that in wild ducks? Virology 1988; 163: 247–250.

8. Danckwerts DK, et al. Biomass consumption by breeding seabirds in the western Indian Ocean: indirect interactions with fisheries and implications for management. ICES Journal of Marine Science 2014; 71: 2589–2598.

9. Jaeger A, et al. Geolocation reveals year-round at-sea distribution and activity of a superabundant tropical seabird, the Sooty Tern Onychoprion fuscatus. Frontiers in Marine Science 2017; 4: 394.

10. Lebarbenchon C, et al. Migratory patterns of two major influenza virus host species on tropical islands. Royal Society Open Science 2023; 10: 230600.

11. Lebarbenchon C, et al. Influenza A virus on oceanic islands: Host and viral diversity in seabirds in the Western Indian Ocean. PLoS Pathogens 2015; 11: e1004925.

12. Clessin A, et al. Circumpolar spread of avian influenza H5N1 to southern Indian Ocean islands. Nature Communications Nature Publishing Group, 2025; 16: 8463.

13. Peacock TP, et al. The global H5N1 influenza panzootic in mammals. Nature 2025; 637: 304–313.

14. Jaquemet S, et al. Comparative foraging ecology and ecological niche of a superabundant tropical seabird: the sooty tern Sterna fuscata in the southwest Indian Ocean. Marine Biology 2008; 155: 505–520.

15. Feare CJ. Ecology of Bird Island, Seychelles. Atoll Research Bulletin 1979; 226: 1–29.

16. Hart LA, et al. Time heals: Boosted breeding seabird populations on restored Cousine Island, Seychelles. African Journal of Ecology 2022; 60: 505–515.

17. Trevail AM, et al. Tracking seabird migration in the tropical Indian Ocean reveals basin-scale conservation need. Current Biology 2023; 33: 5247–5256.

18. Sinclair I, Langrand O. Birds of the Indian Ocean Islands. New Holland Publishers, Limited, 2004.

19. Wilcox BR, et al. Influenza-A viruses in ducks in Northwestern Minnesota: Fine scale spatial and temporal variation in prevalence and subtype diversity. PLoS ONE 2011; 6: e24010.

20. Lebarbenchon C, et al. Comparison of two commercial enzyme-linked immunosorbent assays for detection of influenza A virus antibodies. Journal of Veterinary Diagnostic Investigation 2012; 24: 161–165.

21. R Core Team. R: A Language and Environment for Statistical Computing. Vienna, Austria: R Foundation for Statistical Computing, 2025.

22. Spackman E, et al. Development of a real-time reverse transcriptase PCR assay for type A influenza virus and the avian H5 and H7 hemagglutinin subtypes. Journal of Clinical Microbiology 2002; 40: 3256–3260.

23. Tsukamoto K, et al. Subtyping of avian influenza viruses H1 to H15 on the basis of hemagglutinin genes by PCR assay and molecular determination of pathogenic potential. Journal of Clinical Microbiology 2008; 46: 3048–3055.

24. VanDalen KK, et al. Increased detection of influenza A H16 in the United States. Archives of Virology 2008; 153: 1981–1983.

25. Lee M-S, et al. Identification and subtyping of avian influenza viruses by reverse transcription-PCR. Journal of Virological Methods 2001; 97: 13–22.

26. Qiu B-F, et al. A reverse transcription-PCR for subtyping of the neuraminidase of avian influenza viruses. Journal of Virological Methods 2009; 155: 193–198.

27. Zhang Z, et al. A greedy algorithm for aligning DNA sequences. Journal of Computational Biology 2000; 7: 203–214.

28. Brown JD, et al. Prevalence of antibodies to type A influenza virus in wild avian species using two serologic assays. Journal of Wildlife Diseases 2010; 46: 896–911.

29. Lang AS, et al. Assessing the role of seabirds in the ecology of influenza A viruses. Avian Diseases 2016; 60: 378.

30. Mackenzie JS, et al. Isolation of ortho- and paramyxoviruses from wild birds in Western Australia, and the characterization of novel influenza A viruses. Australian Journal of Experimental Biology and Medical Science 1984; 62: 89–99.

31. Razafimandimby F, Amy M, Le Corre M. Migrating Turnstones use Tromelin Island as an opportunistic stopover site during pre-breeding migration. Wader Study 2025; 131: 219–225.

32. Razafimandimby F, Amy M, Le Corre M. Wintering grounds under protection: population stability and conservation of migrating waders at Europa Island, western Indian Ocean. Ostrich 2025; 96: 258–269.

33. Vally Z, et al. Occasional exposure of native and non-native bird species to avian influenza virus on a remote oceanic island. Submitted.

34. Ramis A, et al. Experimental infection of highly pathogenic avian influenza virus H5N1 in black-headed gulls (Chroicocephalus ridibundus). Veterinary Research 2014; 45: 84.

35. Rijks JM, et al. Mass mortality caused by highly pathogenic influenza A(H5N1) virus in Sandwich terns, the Netherlands, 2022. Emerging Infectious Diseases 2022; 28: 2538– 2542.

36. Abolnik C, et al. Outbreaks of H5N1 high pathogenicity avian influenza in South Africa in 2023 were caused by two distinct sub-genotypes of clade 2.3.4.4b viruses. Viruses 2024; 16: 896.

